# Selection for imperfection: A review of asymmetric genitalia in araneomorph spiders (Araneae: Araneomorphae)

**DOI:** 10.1101/704692

**Authors:** F. Andres Rivera-Quiroz, Menno Schilthuizen, Boopa Petcharad, Jeremy A. Miller

**Affiliations:** Department Biodiversity Discovery group, Naturalis Biodiversity Center, Darwinweg 2, 2333CR Leiden, The Netherlands; Endless Forms Group, Naturalis Biodiversity Center, Darwinweg 2, 2333CR Leiden, The Netherlands; Institute for Biology Leiden (IBL), Leiden University, Sylviusweg 72, 2333BE Leiden, The Netherlands; Faculty of Science and Technology, Thammasat University, Rangsit, Pathum Thani, 12121 Thailand

**Keywords:** Chirality, sexual selection, antisymmetry, Araneae, Synspermiata, Entelegynae, RTA, Liocranidae

## Abstract

Bilateral asymmetry in the genitalia is a rare but widely dispersed phenomenon in the animal tree of life. In arthropods, occurrences vary greatly from one group to another and there seems to be no common explanation for all the independent origins. In spiders, genital asymmetry appears to be especially rare. Few examples have been studied in detail but isolated reports are scattered in the taxonomic literature. Based on a broad literature study, we found several species in thirteen families with evidence of genital asymmetry, mostly expressed only in females. Our review suggests that spider genital asymmetries, although rare, are more common than previously thought and taxonomic descriptions and illustrations are a useful but not entirely reliable tool for studying them. Here we also document thoroughly the case of the liocranid spider *Teutamus politus*. We collected live specimens to observe male-female interactions and document their genital morphology. We consider *T. politus* to be the first known case of directional asymmetry and the first report of developmentally asymmetric male genitals in Entelegynae spiders. Generalities, evolution and categorization of asymmetry in spiders are further discussed.

## Introduction

Genital asymmetry is a trait that has evolved independently several times in many animal groups. Invertebrates show a wide range of genital asymmetries with probably thousands of independent origins. Many, sometimes not mutually exclusive, explanations have been proposed, namely: i) morphological compensation for selected changes in mating position; ii) sexually antagonistic co-evolution; iii) cryptic female choice for asymmetric male genitalia; iv) different functions for the left and right side; v) one-sided reduction to save space and resources; vi) functional constraints: to function properly, the separate parts of the genitalia need to connect in an asymmetric fashion; vii) efficient packing of internal organs in the body cavity [1–4].

Asymmetries are often classified as fluctuating (FA), antisymmetry (AS) or directional (DA) [3,5,6]. This categorization is based on the degree and relative frequencies of the different chiral forms found in a population. FA describes slight asymmetric variation around a symmetrical mean; the appearance of this type of asymmetry is usually related to environmental or developmental constraints [5,7]. AS describes cases where two mirror image forms, dextral and sinistral, are identifiable and within a population, occurring usually in equal or similar proportions [3]. Finally, DA refers to cases where only one asymmetric form is virtually always present [3]; this might be associated with mechanical, behavioral, or functional differentiation and selection of one asymmetrical form of the structures or organs [3,8].

Genital asymmetry, although rare as a whole, is a recurring phenomenon in a few groups of arthropods like mites, crustaceans, opiliones, and very common several insect orders. However, in spiders (Fig. 1a), sexual asymmetries seem to be rather an uncommon exception [1–4,9,10]. In insects, copulatory mechanics and the presence of a single male genital structure located at the posterior end of the abdomen might explain the great incidence of genital asymmetry in this group [1,3,11]. In contrast, spiders have two male copulatory organs derived from a modified pair of leg-like appendages (Fig. 1b). These are normally both used sequentially for sperm transfer during copulation [12]. The presence of these paired structures has been hypothesized to act as an “evolutionary buffer” to the development of genital asymmetry, especially on male genitals [1,3,10].

**Figure 1.**
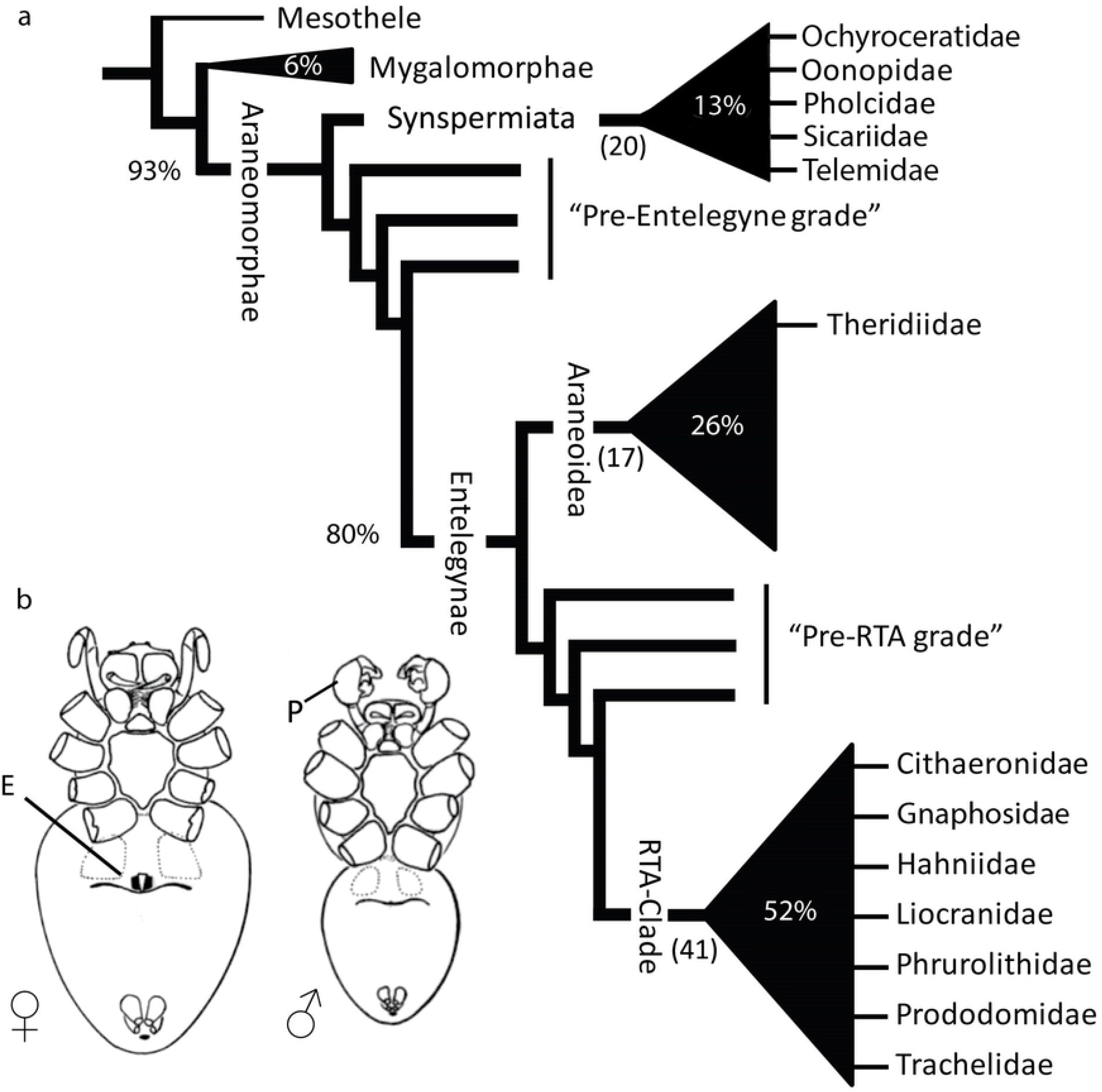
Spider relations and spider genitalia. a) Schematic tree based on a comprehensive spider phylogeny by Wheeler et al. [13]. Number of families per clade are indicated between parentheses; approximate percentage of species per clade relative to the Order Araneae is also given. Family name tags indicate the ones with known asymmetric species. b) Ventral view of spider copulatory organs: ♀ Epigynum (E) and ♂ Pedipalp bulb (P); modified from Foelix [12].

Most cases of asymmetry in spiders have not been studied in detail or even discussed, with the notable exception of pholcids and theridiids [1,3]. Nevertheless, taxonomic illustrations and descriptions give evidence of the existence of this phenomenon in other families. Genital asymmetry has been documented in females, males or both sexes, with seemingly several independent origins in the spider tree of life. All known cases have been reported in two major clades: Synspermiata and Entelegynae that include about 13% and 80% of known spider diversity, respectively (Fig. 1a); within the Entelegynae, asymmetries have been documented in the clades Araneoidea and RTA.

Morphologically, Synspermiata spiders tend to have structurally simpler genitalia than entelegyne spiders in both sexes. Asymmetries in Synspermiata have been properly documented in two families: Pholcidae (Fig. 2 a, h) and Oonopidae (Fig. 2, h); but taxonomic descriptions of some Ochyroceratidae (Fig. 2 b, d), Telemidae (Fig. 2 f) and Sicariidae depict female genital asymmetry too. In Entelegynae, examples appear more scattered with most cases being found in the family Theridiidae (Fig. 3a-c) and some more documented in at least six families of the RTA clade (Fig. 3b, d-h). Explanations for genital asymmetry in spiders are diverse and could include individual variation, natural selection, or sexual selection [1,3,10,14,15].

**Figure 2.**
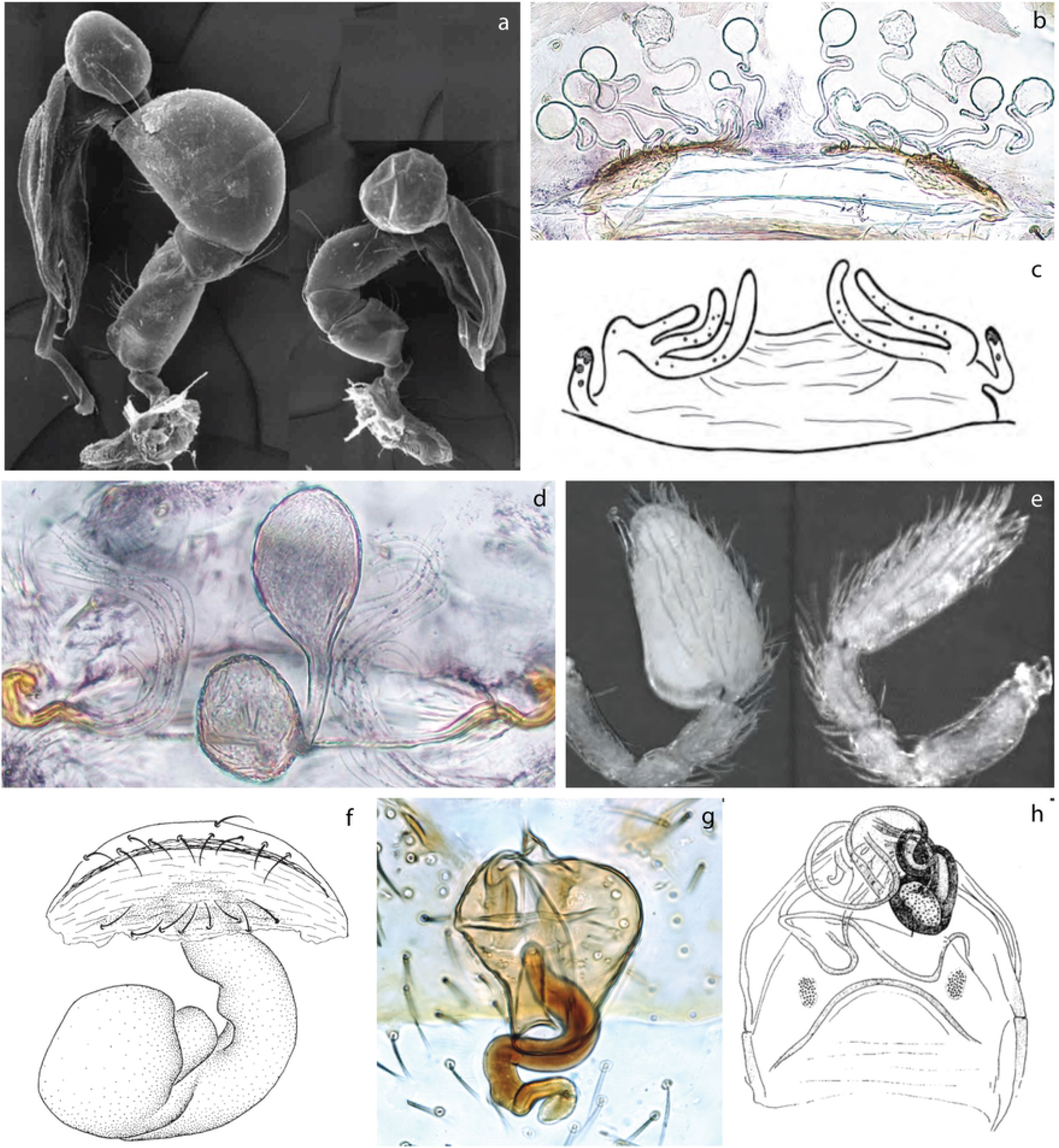
Examples of genital asymmetry in Synspermiata. a, e) male pedipalps, lateral view. b-d, f-h) vulva, dorsal view. a) Pholcidae: *Metagonia mariquitarensis*; modified from Huber [8]. b) Ochyroceratidae: *Althepus naphongensis*; modified from Li *et al.* [16]. c) Sicariidae: *Hexophthalma albospinosa*; modified from Magalhaes and Brescovit [17]. d) Ochyroceratidae: *Speocera cattien*; modified from Tong, *et al.* [18].e) Oonopidae: *Paradysderina righty*; modified from Platnick and Dupérré [19]. f) Telemidae: *Telema exiloculata*; modified from Lin and Li [20]. g) Oonopidae: *Triaeris stenaspis*. h) Pholcidae: *Metagonia delicate*; modified from Huber [21].

**Figure 3.**
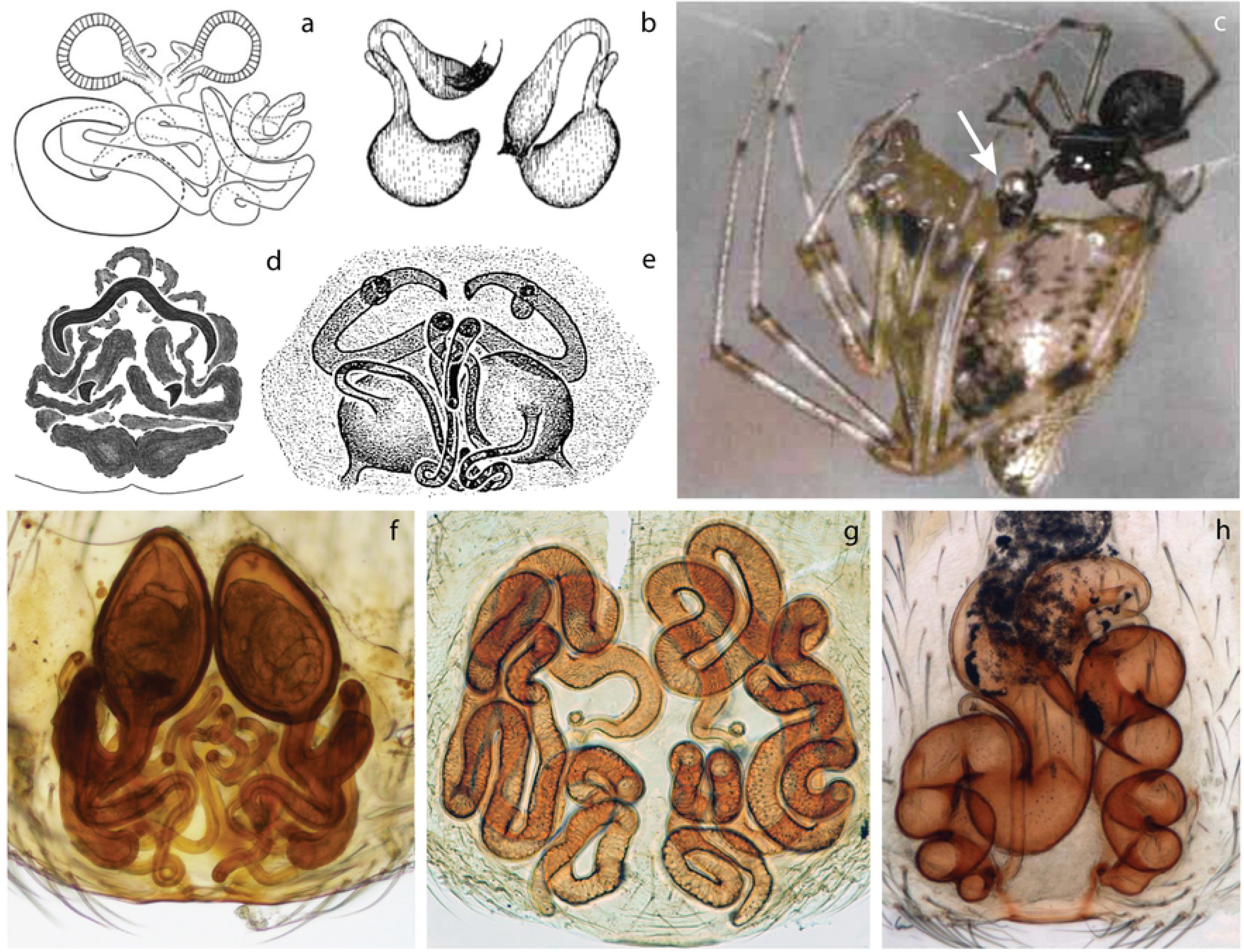
Examples of genital asymmetry in Entelegynae. a, b, d-h) vulva, dorsal view. c) male and female during copulation. a) Theridiidae: *Asygyna coddingtoni*; modified from Agnarsson [22]. b) Phrurolithidae: *Scotinella fratella*,; modified from Dondale and Redner [23]. c) Theridiidae: *Tidarren sisyphoides*. Arrow shows the presence of only one pedipalp; modified from Knoflach [24]. d) Gnaphosidae: *Apopyllus gandarella*; modified from Azevedo *et al.* [25]. e) Hahnnidae: *Neoantistea agilis*; modified from Opell and Beatty [26]. f) Trachelidae: *Trachelas ductonuda*; modified from Rivera-Quiroz and Alvarez-Padilla [27]. g) Liocranidae: *Jacaena mihun*. h) Cithaeronidae: *Cithaeron praedonius*,; modified from Ruiz and Bonaldo [28].

Spider genital asymmetry can be classified as follows: Fluctuating asymmetry (FA) is probably the most common type and has been properly documented in some Lycosidae [29–32], Pholcidae [33], and Oxyopidae [10,34]. Other examples of seemingly asymmetric structures like the pedipalps of the one known specimen of *Pimoa petita* [35] or the numerous documented anomalies and deformities [36–39] might easily be explained by developmental malformations (Fig. 4).

**Figure 4.**
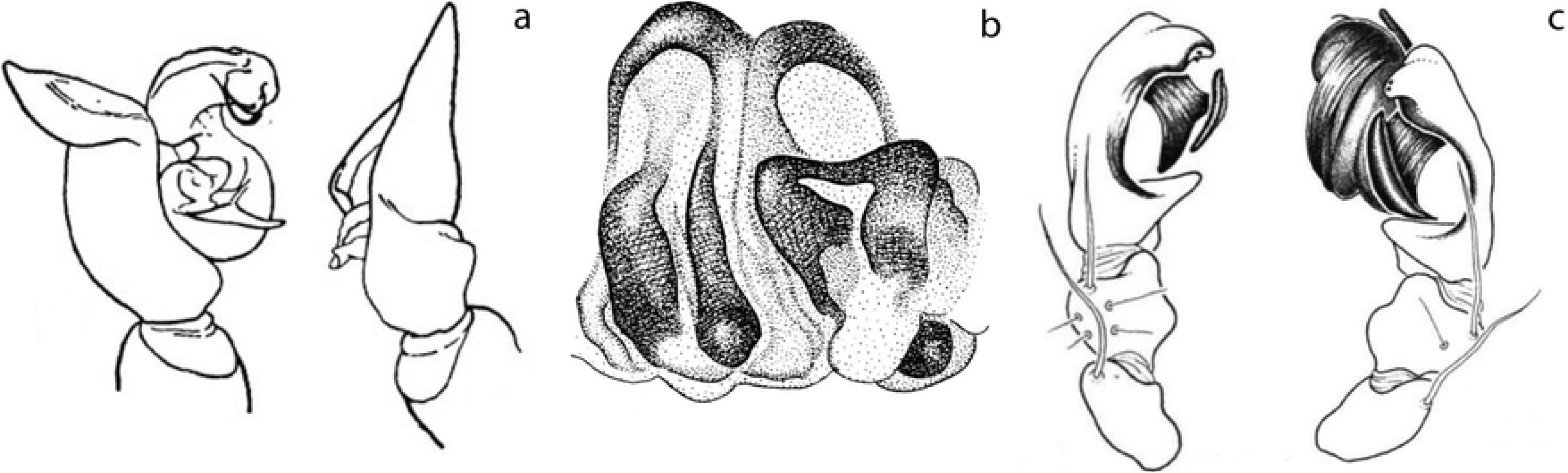
Examples of genital malformation in spiders. a,c) male pedipalps, posterior-lateral view. b) vulva, ventral view. a) *Lycosa ammophila*; modified from Kaston [37]. b) *Pardosa sagei*; modified from Kaston [37]. c) *Pimoa petita*; modified from Hormiga [35].

Antisymmetry (AS) is the second most common form of asymmetry in spiders and has been documented in three genera of the Theridiidae (*Asygyna, Echinotheridion*, and *Tidarren*) (Fig. 3a, c) [22,40,41]; one genus of Pholcidae (*Metagonia*) (Fig. 2a, h) [21]; one genus of Phrurolithidae (*Scotinella*) (Fig. 3b) [42] and scattered cases such as in Trachelidae (Fig. 3f) [27,43,44], Cithaeronidae (Fig. 3h) [45] and other RTA families. Directional asymmetry (DA) is the rarest type and, until now, it had only been reported in the pholcid *Metagonia mariguitarensis* (Fig 2h) [8]; DA has also been implied some descriptions within the Oonopidae (Fig 2e) [19,46], and in the liocranid *Teutamus politus* female genitalia [47]. All of these, other isolated reports, and scattered descriptions and illustrations suggest that genital asymmetries in spiders have originated independently several times and their study might give better insights into how and when this phenomenon has evolved and the selective mechanisms behind it.

A particularly interesting example are the Liocranidae where two different types of asymmetry are present [47–49]. For example, *Jacaena mihun* (Fig. 3g) shows no external chirality, but internally the asymmetric copulation ducts are highly variable among individuals. Another example, *Teutamus politus* (Figs 5–7), shows external asymmetry in the female genitalia with both copulatory openings fused together in one atrium placed on the left side of the epigyne (see Deeleman-Reinhold [47]: fig 800, 801). Deeleman-Reinhold [47] mentioned female asymmetry as a diagnostic character for this species and noted that in all six of the specimens available for examination, the atrium is located in the left side. A revision of the genus *Teutamus* [48] also included asymmetry in the female genitalia as a diagnostic character for *T. politus*, and expanded the sample of specimens examined; asymmetry in male pedipalp was not reported in either of these cases.

**Figure 5.**
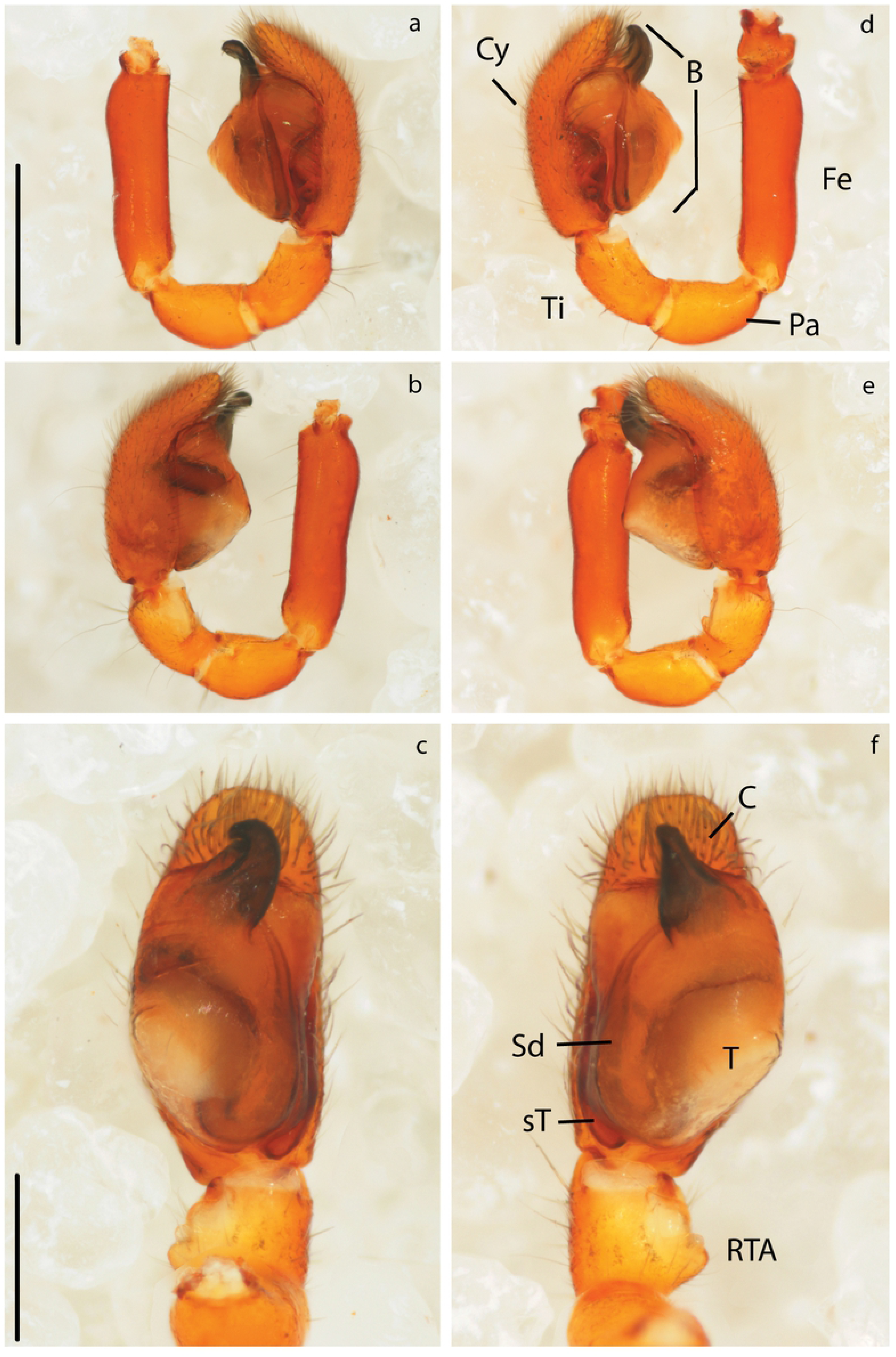
Asymmetric male genitalia of *Teutamus politus*. Right pedipalp: a) prolateral view. b) retrolateral view. c) ventral view. Left pedipalp: d) prolateral view. e) retrolateral view. f) ventral view. Scale bars: **a**, **b**, **d**, **e** = 0.5 mm. **c**, **f** = 0.25 mm.

Here we present a general review of genital asymmetries in spider literature, grouping them in previously described categories of genital asymmetry and discussing the existence of a new category of female genital asymmetry (here called Chaotic Asymmetry). We also analyzed the specific case of the species *Teutamus politus* by collecting new specimens in Thailand and documenting male and female genitalia using diverse morphological methods. This gives evidence of the first cases of both directional asymmetry in males and females, and developmental male genital asymmetry in Entelegynae spiders.

## Material and Methods

### Literature review

We performed an informal search in taxonomic literature of several Synspermiata and Entelegyne families. Selection of publications was initially based on reported cases in literature [1,3,8,10,11] and then expanded depending on the occurrences found within each family. We did not contemplate individual cases of clear FA but this type of asymmetry is included in our discussion. We considered *T. politus* as a good model for testing basic hypotheses on genital asymmetry because of the clear external and internal morphology of female genitalia and Deeleman-Reinhold’s [47] note suggesting this could be a case of DA. Furthermore, we hypothesized that morphological or behavioral compensation for female genital asymmetry could be found in the male.

We considered male asymmetry as those cases that result in clear morphological differences between right and left pedipalp regardless of having a developmental or behavioral origin. Based on this, we also considered the pedipalp amputation that males of *Echinoitheridion* and *Tidarren* perform on themselves in our review; especially since the asymmetry has clear adaptive and evolutionary implications [14,41,50–52].

### Fieldwork

We selected study sites and collecting dates based on the relative numbers of collected adult specimens of *T. politus* mentioned in literature [47,48]. Fieldwork was carried out in Thailand between July 29^th^ and August 12^th^ 2018; here we sampled 12 sites in total: eight in Phuket Island and four more in Krabi Province. We attempted to cover a variety of vegetation types ranging from relatively well preserved mixed forests to rubber and oil palm plantations. In each site we processed leaf litter using Winkler extractors and direct collecting on ground, among leaf litter and under rocks and logs. Hand collected specimens were kept alive in individual tubes. Winkler specimens were collected in a mixture of propylene glycol and 96% ethanol. All the specimens have been deposited in the collection of the Naturalis Biodiversity Center, Leiden, The Netherlands.

### Behavioral observations

Live specimens were kept individually in clean 15ml Falcon tubes and fed with termites every two days. Seventeen males and 19 females were selected and assigned unique numbers. Couples were formed preferably with specimens from the same locality. Spiders were placed in a Petri dish (diameter 5 cm, height 1 cm); each dish was divided by a paper wall with a small opening so spiders could roam freely but flee in case of aggression. Each couple was kept in the dish under constant observation for a period of about three hours. After observations, all specimens were sacrificed and stored in 96% ethanol.

### Morphological methods

Somatic characters and male sexual structures were photographed using a Leica MI6SC Stereomicroscope equipped with a Nikon DS-Ri2 camera. Female genitalia were dissected, digested using a pancreatine solution [53], cleared with methyl salicylate. Observations were made using semi-permanent slide preparations [54] in a Leica DM 2500 microscope with the same camera as above. Male genitals were expanded using 10% KOH and distilled water in three 3 min. cycles leaving the pedipalps in distilled water overnight to stabilize them for photography. Female epigyna and male pedipalps were prepared for SEM and mounted following Alvarez-Padilla and Hormiga [53] SEM images were obtained using a JEOL JSM-6480LV electron microscope.

The following abbreviations are used in the text and figures: **Female genitalia**: A, atrium; CD, copulatory ducts; CO, copulatory openings; Fd, fertilization ducts; Sa, secretory ampullae (*sensu* Dankittipakul, Tavano, and Singtripop [48]); S, spermatheca. **Male genitalia**: B, male pedipalp bulb; Cy, cymbium; C, pedipalp conductor; E, embolus; Fe, femur; H, basal hematodocha; Pa, patella; RTA, tibia retro lateral apophysis; Sd, sperm duct; sT, sub tegulum; T, tegulum; Ti, tibia.

## Results

### Literature review

We reviewed publications that directly focus on genital asymmetry as well as taxonomic literature that tangentially describe or illustrate asymmetrical morphology. We found ca. 150 species across thirteen spider families with indications of asymmetric genitalia (Table 1) representing less than 0.3% of all spider species World Spider Catalog [55]; and about 13.5% of all the currently valid species in the genera reviewed for this study. Synspermiata has at least five families (Ochyroceratidae, Oonopidae, Pholcidae, Sicariidae and Telemidae) where some kind of asymmetry has evolved accounting for ca. 90 species (Table 1). Asymmetry was found in both female and male genitalia; female asymmetry is more frequent, being found in at least five oonopid, three sicariid, two pholcid and two ochyroceratid genera. In addition, most genera in the Telemidae have evolved a single sac-like seminal receptacle; some species show seemingly asymmetric modifications of this sac, leaning and sometimes spiraling to one side. However, intraspecific variation has not been documented. Male asymmetry is less common, being found in three oonopid and two pholcid genera, and ambiguously suggested for two ochyroceratid species [18,56]. Nevertheless, it is prevalent in *Escaphyella* and *Paradysderina*, where about 20 species show asymmetric male pedipalps (2 e).

**Table 1.**
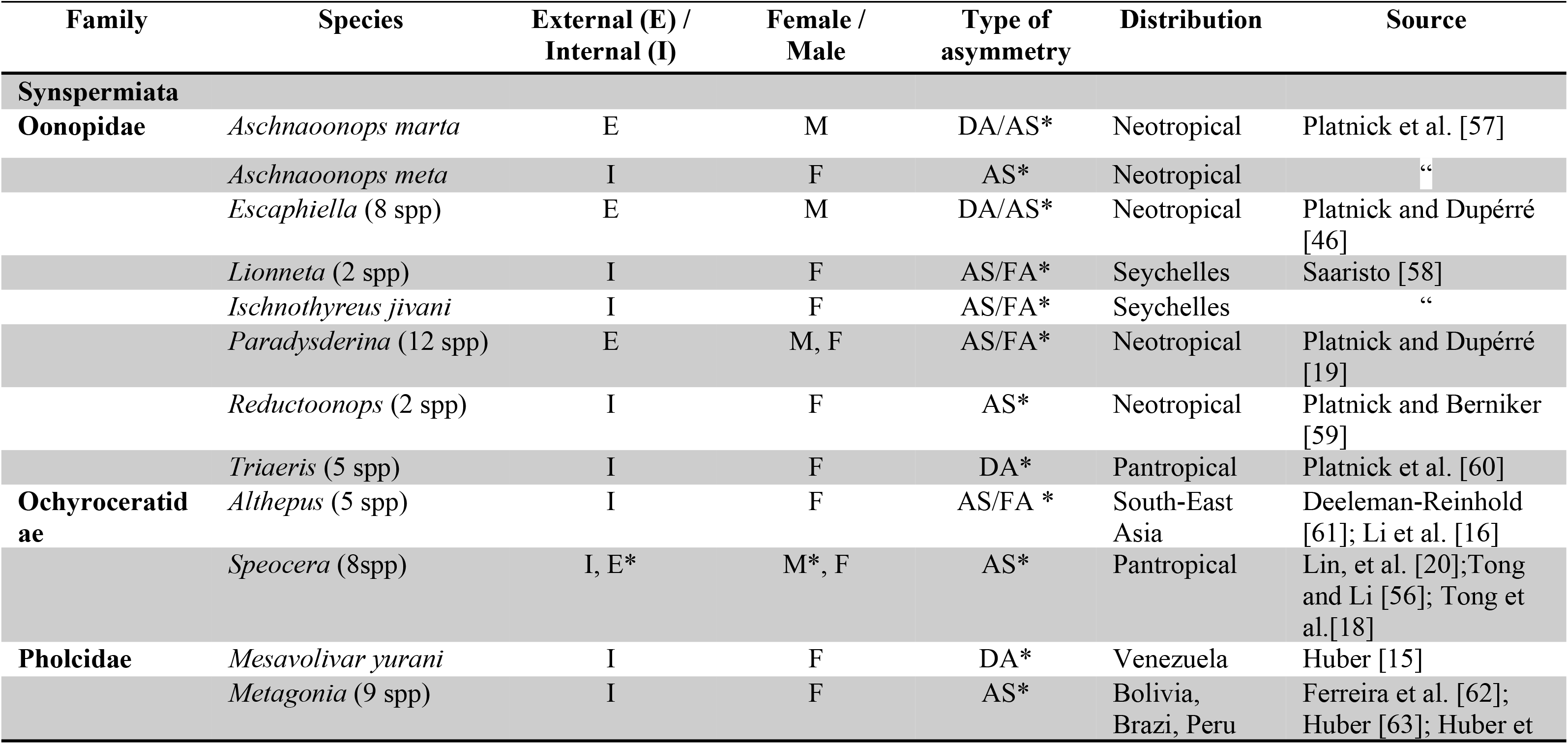
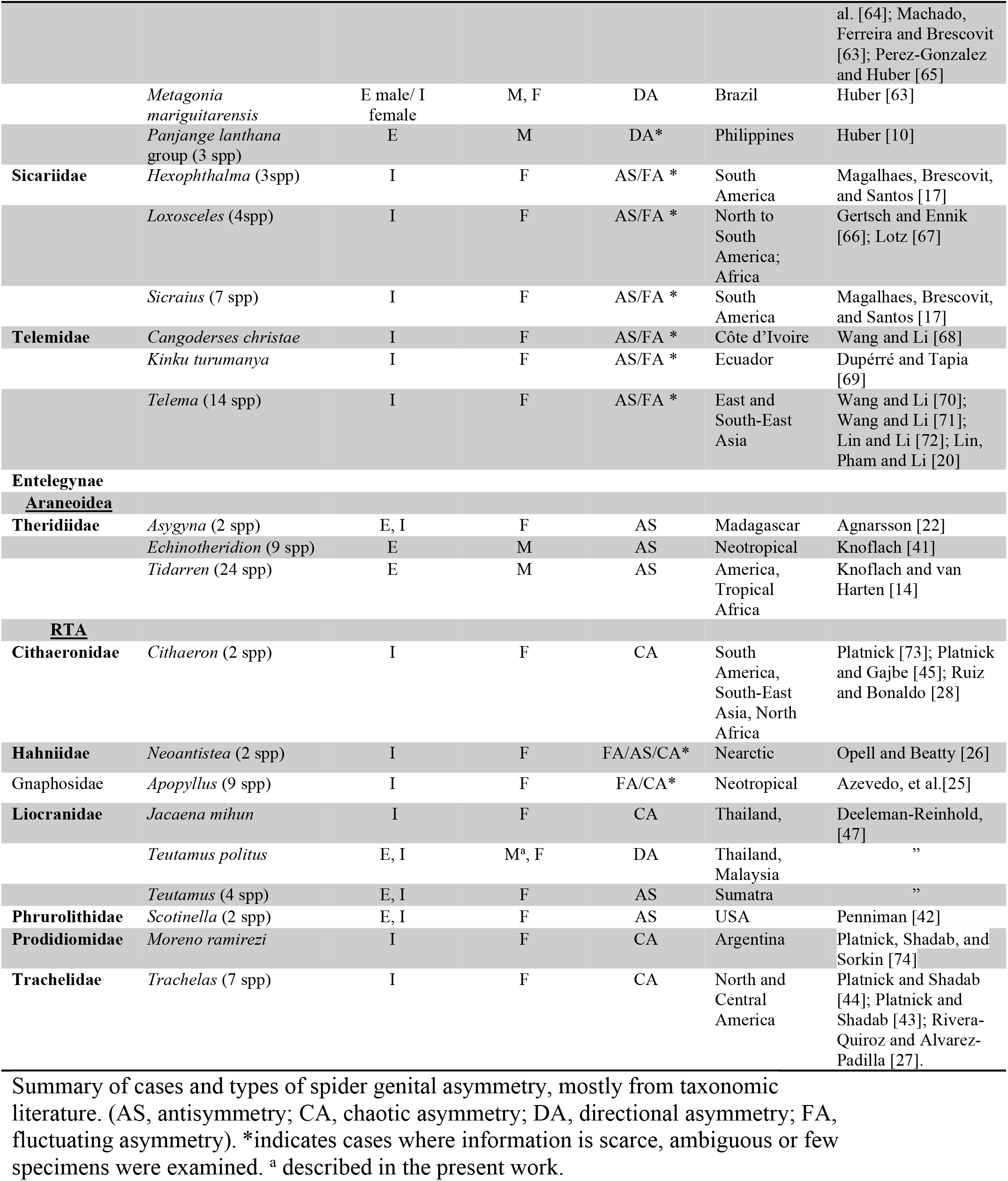
Spider taxa with genital asymmetry reports in literature.

In Entelegynae, more than 60 species in eight families show genital asymmetry. Almost half of the cases were found in the Theridiidae with ca. 35 species in three genera (*Asygyna*, *Echinotheridion*, and *Tidarren*). The rest are scattered among seven families in the RTA clade (Cithaeronidae, Hahniidae, Gnaphosidae, Liocranidae, Phrurolithidae, Prodidiomidae, Trachelidae) (Table 1). Most genital asymmetry reports in Entelegynae include only female genitalia. Female internal asymmetry was the most common, showing a wide range of variation on spermathecae and copulatory ducts (Fig. 3d-h). Female external asymmetry was only found in *Asygyna* (Fig. 3a), *Scotinella* (Fig. 3b) and *Teutamus* (Fig. 7a,d). Male genital asymmetry in Entelegynae had only been reported in the theridiid *Echinotheridion* and *Tidarren* (Fig. 3c); these two genera exemplify a unique behavior that results in genital mutilation; however, normal developmental asymmetry, rather than behaviorally induced, had never been described in Entelegynae literature before this work.

Most male asymmetries in literature appear to be AS with the exception of the DA in *Metagonia mariguitarensis*, two species of *Escaphiella* and the newly described pedipalps of *Teutamus. politus*. Three species of the *Panjange lanthana* group, and several more of *Aschnaoonops*, *Escaphiella*, and *Paradysderyna* might also be DA but only a few specimens have been examined. Female genital asymmetry most of the times involves only internal structures such as ducts, bursa, and spermatheca. Both AS (Fig. 3a-b) and CA (Fig. 3d-h) are relatively common. External asymmetry is not usual and had only been described in *Asygyna*, Theridiidae (Fig. 3a) and *Scotinella*, Phrurolithidae (Fig. 3b) (apparently AS); and *Teutamus*, Liocranidae (Fig. 7) (DA and apparent AS).

A total of 60 female and 35 male specimens were collected as a result of our fieldwork in Thailand. External female genitalia and male pedipalps were observed and compared for all specimens. Four females and five males had their genitals dissected and prepared for detailed examination.

### Male genital morphology

All pedipalp segments with the exception of the bulb (B) seem to be completely symmetrical. Bulbs show at least three clear asymmetries between the right and left sides: i) left B is slightly wider than the right one (Fig. 5b,e; c, f; 6a); ii), left side has a flatter and wider tegulum (T) (Fig. 5f) projected anteriorly in retrolateral view (Fig 5e); and iii) the left conductor (C) is conical and straight (Fig. 6b), slightly pointing towards the cymbium (Cy) in lateral view (Fig. 5d, f); while the right C is flattened, hook-shaped (Fig. 6c) and pointing away of the Cy in lateral view (Fig. 5a, c). There is no apparent difference in the length and shape of the emboli (E) or the spermatic ducts (Sd). This suggests that the asymmetry might not be linked to functional distinction of left and right pedipalp.

**Figure 6.**
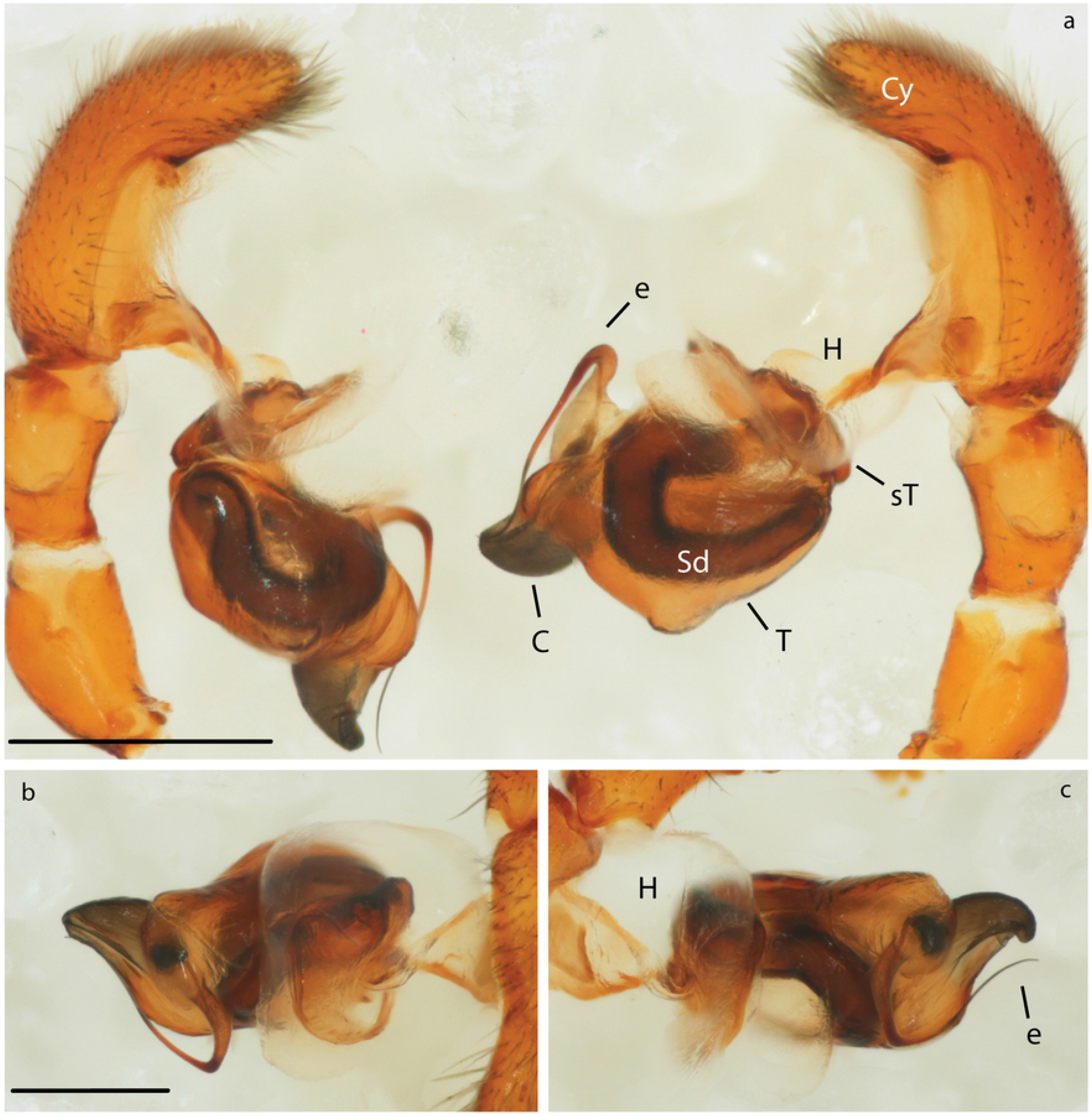
Expanded asymmetric male genitalia of *Teutamus politus*. a) comparative retrolateral view. b) left pedipalp prolateral view. c) right pedipalp prolateral view. Scale bars: **a** = 0.5 mm. **b**, **c** = 0.25 mm.

### Female genital morphology

Externally, the epigynal plate is flattened and fused to the ventral scutum (Fig. 7a). Copulatory openings (CO) are placed close together, forming an atrium facing the left side of the venter and located anteriorly to the bean-shaped spermatheca (Fig. 7a-c). Left spermatheca is slightly shorter than right one (Fig. 7c). Copulatory ducts (CD) are equally long. Right CD anterior to the right spermatheca, left CD located in between both spermathecae (Fig. 7c, e). Asymmetric attachment of CD to spermathecae with the right being anterior to that of the left one (Fig. 7b, c). Both CD have secretory ampullae (Sa) close to their middle portion (Fig. 7b, c). Fertilization ducts (Fd) short and simple, originating from the posterior end of the spermatheca and pointing in the same direction (Fig. 7 e). Despite the clear difference in shape, there is no morphological evidence that suggests functional differentiation between right and left structures.

**Figure 7.**
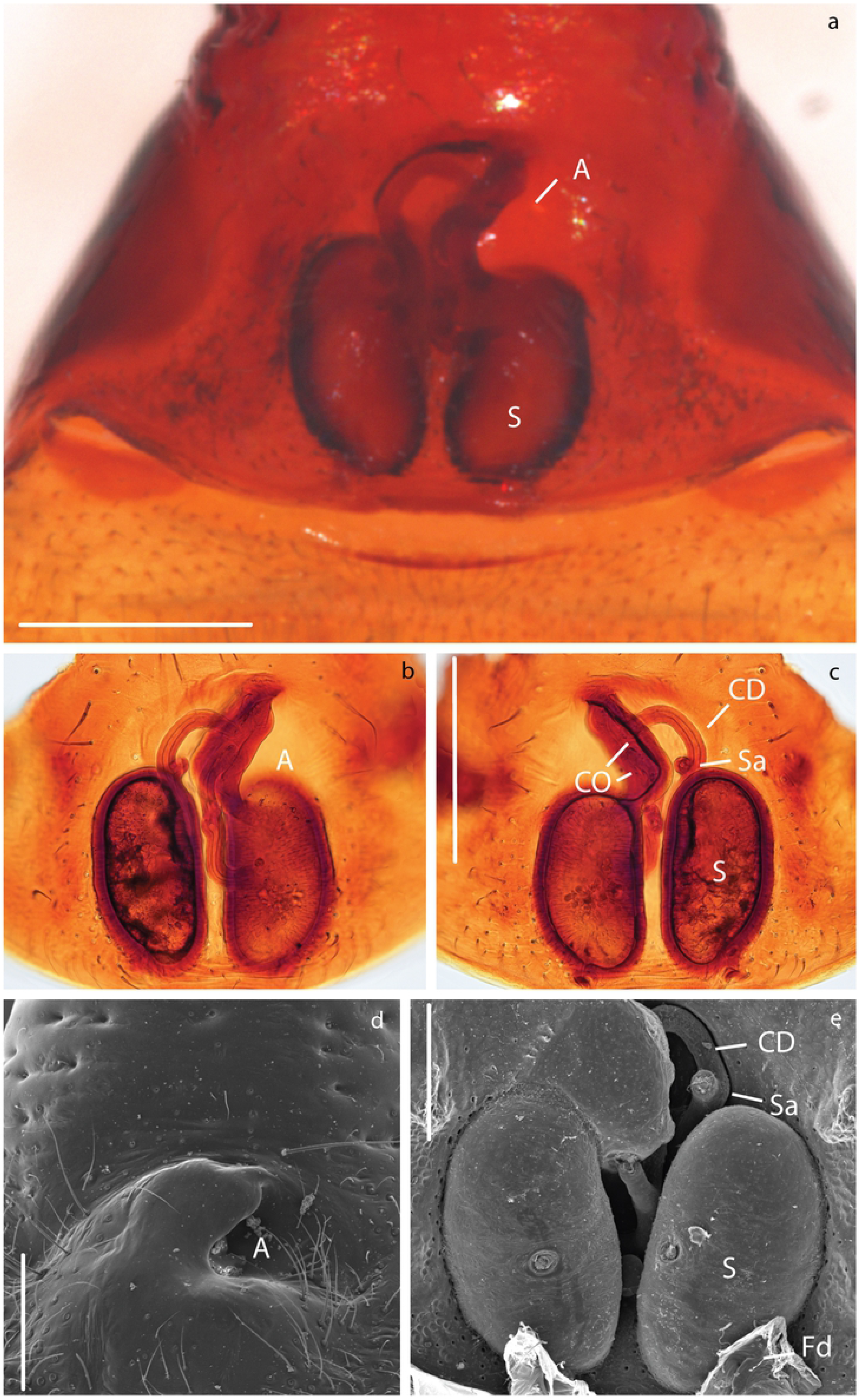
Asymmetric female genitalia of *Teutamus politus*. a) epigynum ventral view. b) dissected and cleared vulva ventral view. c) same, dorsal view. d) vulva, ventral view, SEM. e) same, dorsal view. Scale bars: **a**, **b**, **c** = 0.25 mm. **d** = 150 um. **e =** 100 um

### Behavioral observations

A total of 25 different couples were tested. Initially couples were formed with males and females from the same collection site. Males were more difficult to keep alive than females with most males dying within three days of collection. Due to this, males and females from different sites were also coupled. There were no successful observations of either courtship or mating. Spiders preferred to explore the dish or stand still and, whenever they got too close, they usually avoided each other. In general, interactions between females and males were brief and non-aggressive. Four females laid egg sacs in the Falcon tubes.

## Discussion

### Literature review

Taxonomic literature is the biggest repository of primary descriptive data on the world’s biodiversity. However, illustrations and description are difficult to interpret and might be influenced by the number of studied specimens, state of preservation, preparation artifacts and even illustration techniques. As an example, the species *Cithaeron indicus* shows clear asymmetric female genitalia in its original description [45] but appears symmetrical in a later publication [75] (Fig. 8). Illustrators sometimes avoid introducing variation by drawing one half of a given structure and then tracing the other side based on it. This might simplify understanding and drawing some structures but could also lead to overlooking important information in the illustration process. Similar biases have been observed in some species of *Trachelas* [43,44] and could be present elsewhere. As pointed out by Huber and Nuñeza [10], preparation artifacts might also play a role in the identification and interpretation of asymmetric structures. Weakly sclerotized internal genitalia (as that typically found in non-Entelegynae spiders) are often prone to create artifacts during specimen preparation and an interpretation without sufficient knowledge of intraspecific variation might be misleading. Entelegyne spiders tend to have more heavily sclerotized bodies being less sensible during the preparation process and allowing a more robust interpretation of their genital morphology.

**Figure 8.**
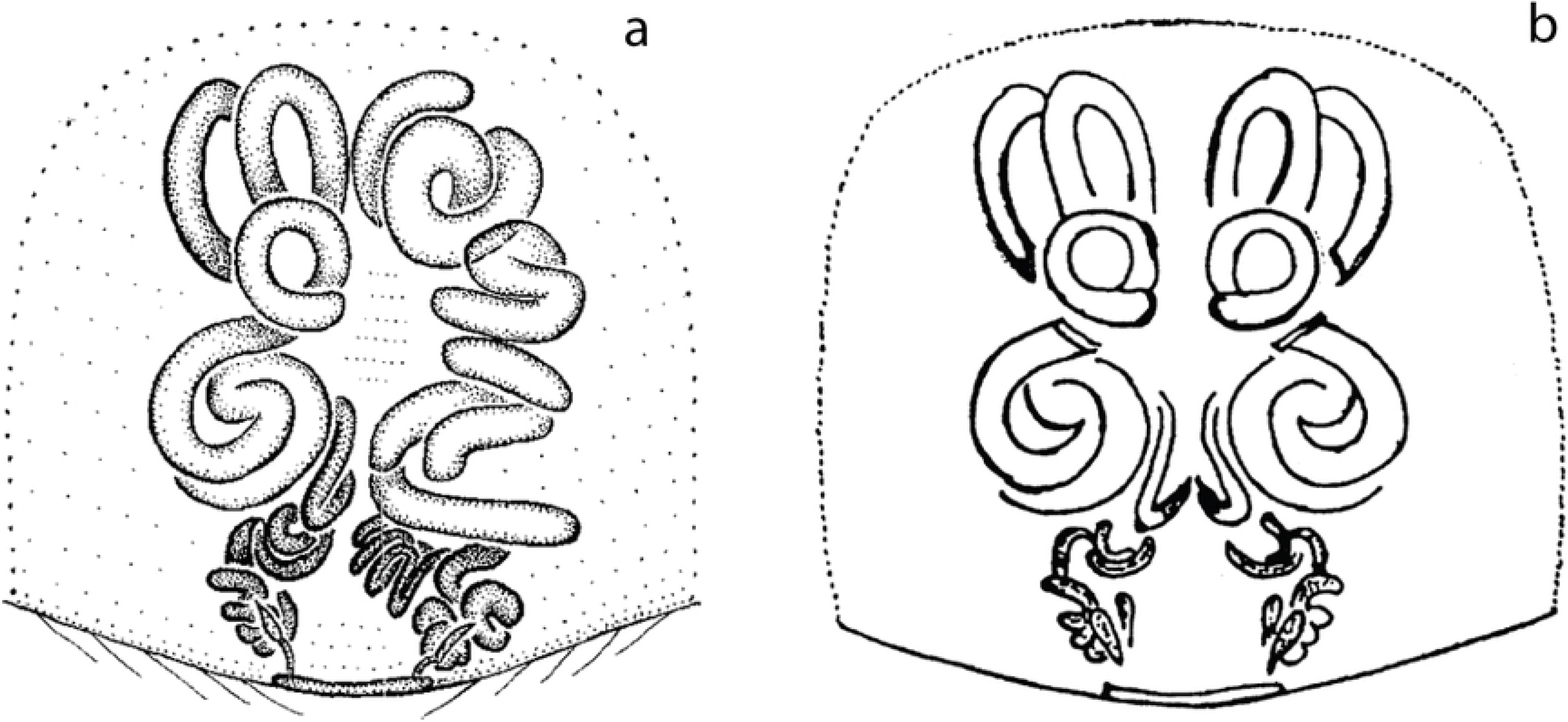
Example of illustration bias. Vulva, ventral view. a) *Cithaeron indicus*; modified Platnick and Gajbe [45]. b) Same; modified from Gajbe [75].

Descriptions of male spider genitalia are also subject to preparation artifacts or methodological biases. Male genitalia preparation and examination is usually done by dissecting, studying and illustrating only one pedipalp. Although this is a very efficient approach and does not represent a problem on most occasions, some cases of asymmetric genitalia might go unnoticed. This has resulted in a more difficult assessment of male asymmetry; as an example, *Metagonia mariguitarensis* was believed to be the only species with male genital asymmetry [8]. However, DA in males of *T. politus* had never been discovered before, apparently because the right male pedipalp had simply been overlooked in previous descriptions. Similarly two *Speocera* species [18,56] have their male pedipalps ambiguously described as “asymmetric” but no more details were given.

In comparison, recent revisionary studies on the oonopid genera *Aschnaoonops*, *Escaphiella*, *Paradysderyna* and *Reductonoops* [19,46,57,59] took special care in comparing the right and left male pedipalps revealing many more cases of genital asymmetry. In all of these, male pedipalps show clear differences in bulb development and embolus shape between right and left (Fig. 2 e). In at least two species of *Escaphiella* [46] enough specimens have been examined to suggest that asymmetry in these species is directional.

### Evolutionary trends of genital asymmetry

We found evidence of ca. 150 cases of asymmetry in spider genitals in thirteen different families. In previous broad-scope reviews, only some examples in Pholcidae and Theridiidae had been taken in account. Reports on insects suggest that genital asymmetry rarely appears isolated and is usually a shared trait between closely related species [3,4,76]. Here, we found some similar patterns with several species within a genus showing at least one type of genital asymmetry. This pattern is more common in the Synespermiata, but was also observed in Entelegynae (Table 1). Although the known number of cases and families with asymmetrical genitalia has increased significantly, this still represents less than 0.3% of all known spider species. The low incidence of genital asymmetry in spiders has been mainly explained by the presence of two sperm transfer structures in the male [1,3]. Huber, Sinclair, and Schmitt [1] remark that in comparison to insects, most spider asymmetry originates in females instead of males, and most insect asymmetry originates as DA, while most or all spider asymmetry originates as AS. Many examples support the first claim, which also fits a cryptic female choice hypothesis [9]. Nevertheless, we found numerous “new” examples of male asymmetry hidden in taxonomic literature (Table 1), highlighting the many cases in the Oonopidae where male asymmetry has apparently not coincided with modified female genitalia. As for the second claim, we found that DA might not be as rare as previously thought. Examples of DA include two confirmed cases in *Metagonia* [8] and *Teutamus*, both involving male and female genitalia; two more in *Escaphiella* [46], that include only male pedipalp; and some more in *Panjange* [10], *Mesovolivar* [15], and three Oonopidae genera that suggest asymmetry directionality but are not conclusive.

Many spider asymmetries seem to fit in the AS category, although only a handful have been evaluated for the appearance of right or left-sided asymmetries within a sample as in Phrurolithidae and Theridiidae [22]. Also, we found some cases in which female copulatory ducts are long, coiled and entangled in a way that does not fit any of the three known types of asymmetry. We called this chaotic asymmetry (CA) because the great variation between individuals of the same species does not allow distinguishing either a dextral or a sinistral form.

Other cases difficult to assess are: the reduction of spermathecae to a single receptacle, as seen in some oonopids [35,51,52], pholcids [62,63,65], and telemids [20,68–72] (Fig. 2f, g); and the presence of odd numbered spermathecae in some sicariids [17,66,67,77] and ochyroceratids [16,61,78] (Fig. 2 b, c). Both phenomena can sometimes generate a seemingly asymmetric morphology. Although good illustrations and photographs of these are available in literature (e.g. figs. 20: Magalhaes, Brescovit, and Santos [17]; figs 14 and 19: Li *et al.* [16]; fig. 8: Lin, Pham and Li [20]; fig. 7: Wang and Li [70]) only some cases in the Sicariidae [17,66] have reported intraspecific variation.

A correct interpretation of the type of asymmetry based only on the available literature is complicated. Many cases describe single specimens or small samples and do not include enough information to assess the character variation within the species. This is a key piece of information since the proportions of forms within the population are crucial to distinguish the type of asymmetry and the evolutionary mechanisms behind it. Here we include examples that, to the best of our knowledge, fit the definition of each type of genital asymmetry and give hypotheses that could explain their origin.

### Fluctuating Asymmetry (FA)

This kind of asymmetry is defined by van Valen [5] as “the inability of organisms to develop in precisely determined paths”. In other words, FA describe random morphological fluctuations around a symmetric mean [3,5,33,79,80]. FA incidence, relation to environmental factors, and its influence within populations has been studied on some Lycosidae and Pholcidae [17–21]. Here we found that some cases, like the hahniid *Neoanthistea*, some oonopid and telemid genera (mentioned as FA* in Table 1), and other “malformed” specimens in literature might be cases of FA. Similarly, the great intraspecific variation observed in the female genitalia of some sicariids [17,66], range from asymmetries in number, size and shape of spermathecae to almost symmetric structures. This suggests that asymmetries in this family and similar cases in the ochyroceratid *Althepus* [16,61] might be fluctuating. Nevertheless, most FA asymmetries require more specimens and careful examination to determine better the nature of the observed phenotypes. A few species that show AS (*Scotinella britcheri* and *S. fratella*) and all species with CA (*Cithaeron praedonius*, *Jacaena mihun*, among others) show great morphological variation of female internal genitalia within the population; however, these variations are never around a symmetric mean and thus we do not consider them to be fluctuating.

### Antisymmetry (AS)

We found this type of asymmetry in two different families and at least four genera; notably, all known species of *Echinotheridion* and *Tidarren* share this trait. This specific case of AS induced by an uncommon genital automutilation behavior may also be the best studied and understood. In these theridiid genera, male spiders show no preference for either left or right pedipalp self-emasculation, and no selection of right or left form by females has been observed. Likewise, females of *Scotinella britcheri* and *S. fratella* show two basic forms with some range of variation in-between but no significant predominance within the studied population [42]; *Asygina huberi* and *A. coddingtoni* [22], and probably some *Teutamus* species like *T. brachiatus*, *T. poggi*, and others (as illustrated by Dankittipakul, Tavano, and Singtripop [48]) also fit this AS model. Likewise, some asymmetric species in the Ochyroceratidae, Oonopidae, Pholcidae, and Telemidae could show AS. However, larger numbers of specimens are needed to identify the proportion of forms and type of asymmetry.

Palmer (1996) divided asymmetry as genetic (larval) and external (post-larval) depending on the developmental stage where it is originated. In spiders, genital development is only apparent after the last molt. Therefore, the exact moment where AS appears, especially in females, is difficult to interpret. Evidence on snails [81], crustaceans [6], and insects [6,76,82] suggest genetic AS to be an evolutionarily unstable or transitional state between symmetry and DA, or even a reversal phase from DA [3,6]. Similarly, a genetic assimilation process of external AS could ultimately lead to DA [3,6,83]. We consider the few confirmed spider AS to represent both types of asymmetry: genetic in *Asygyna* and *Scotinella*, and external in *Echinotheridion* and *Tidarren*.

Another interesting observation is the sex biased incidence of AS. This seems to be also the case in some insect groups like Odonata, Ortopthera, Mantodea, and others [1,2,76]. In *Asygyna* and *Scotinella*, asymmetry has only been reported in females; while the theridiids *Echinotheridion* and *Tidarren* only show asymmetry on male pedipalps. Appearance of AS in *Asygyna* and *Scotinella* might be related to an intrasexual competition between females; while, AS in *Echinotheridion* and *Tidarren* is considered an example of antagonistic co-evolution derived from the extreme size dimorphism between sexes [14,41,50–52]. Neither mechanical, behavioral nor functional differentiation between chiral forms has been reported in the cases above.

### Chaotic asymmetry (CA)

This new category of asymmetry does not fit the definition of any of the three traditional types. Females usually develop long and convoluted copulation ducts where the great variation between specimens does not allow a clear distinction between a dextral and sinistral form. All known examples of this type of asymmetry are found in the Entelegynae clade. Platnick [73] mentioned for *Cithaeron praedonius* (Cithaeronidae): “No two females show identical patterns of epigynal duct coiling; for that matter, no individual specimen shows identical coiling of the ducts of the right and left sides”. Similar morphological variation (Fig. 3d-h) has been observed in some species of the following genera: *Apopyllus* (Gnaphosidae) [25], *Neoantistea* (Hahniidae) [26], *Moreno* (Prodidiomidae) [74], *Jacaena*, (Liocranidae) [49] and *Trachelas* (Trachelidae) [27,43,44].

The origin of these internal genital modifications has not been investigated and its relation to a functional differentiation between sides or packing of other internal organs cannot be ruled out. We hypothesize that the development of this kind of asymmetry is related to complexity in internal female genitalia and this could explain the absence of examples in the genitalicly simple Synspermiata. The absence of a clear right/left pattern and great variation between individuals suggest that copulatory duct shape is not under a strict selection. This might be related to a simplification in pedipalp sclerite complexity and embolus length (as seen in *Trachelas*, *Jacaena* and *Moreno*). In contrast, some *Apopyllus* males have fairly complex male genitals with an extremely long embolus that usually coils around the bulb. Female ducts show slight asymmetries between right and left sides and authors mention internal variation between conspecific females. This genus also shows intraspecific variation in the RTA and external genitalia and it is hypothesized to be an instance of male-female coevolution [25]. The cases of *Cithaeron indicus, Moreno ramirezi* and both *Neoantistea* species are doubtful; in the former, the male is not known, and in *Moreno* and *Neoantistea*, species were described based on just one female or variation was not documented; the observed asymmetry could be fluctuating, antisymmetric, a developmental abnormality or even an artifact of preparation.

If pedipalp bulb sclerite reduction is related to the appearance of CA, the question would be why is it so rare? Within Entelegynae, several groups have reduced male pedipalp complexity; however, CA has not evolved nearly as many times. This might be explained by the evolution of long and convoluted CD prior to male sclerite reduction which would, hypothetically, reduce selective pressure on the female copulatory ducts.

### Directional asymmetry (DA)

In insects, DA is the most common type of asymmetry [1,2]; however, in spiders, DA seems to be quite rare. So far, in Synespermiata, only the pholcid *Metagonia mariguitarensis* [8] had been confirmed as DA; however, there are reports of consistent one-sided asymmetries in other members of this clade. In *Escaphiella gertschi* and *E. itys*, all examined males have developmental differences between right and left pedipalp [46]. More asymmetric male pedipalps have been described for three *Panjange* species of the *lanthana* group [10], *Aschnaoonops marta* [57], at least six species of *Paradysderina* [19], and several species of *Escaphiella* [46]. Likewise, female internal genitalia of *Mesovolivar yurani* [15]; and some species of *Paradysderina* [19], *Reductoonops* [59] and *Triaeris* [60] show asymmetries that seem to be consistent within their samples; nevertheless, the number of specimens examined in many cases is too small to confirm directionality.

The story seems to be different for Entelegynae spiders where more complex development of genitals might inhibit the evolution of directional asymmetry. Although implicit in the description of *Teutamus politus* female genitalia by Deeleman-Reinhold (2001), the present study is the first report of DA in the entelegyne clade. *Teutamus politus* is also the first example of developmental male genital asymmetry in the Entelegynae. Previously, male asymmetry in this clade was only known from teratogenic specimens and the unique AS phenotype created by self-emasculation in *Tidarren* and *Echinotheridion*.

Putative cases of male DA in *Escaphiella* and other oonopids may not be related to modifications in female genitalia [46,57] but to functional segregation of the right and left pedipalps. Similarly, the genus *Triaeris* has many cases of female genital asymmetry that have not been linked to male pedipalp modifications. In fact, some species of this genus are believed to be parthenogenetic [60]. In contrast, directional genital asymmetries in *M*. *mariguitarensis* and *T. politus* have been found in both sexes, which might indicate that selection by female choice is the underlying cause. In these species, males would have to change morphologically or modify mating positions to be able to have successful copulation. Morphological modifications in both sexes have been confirmed for *M. mariguitarensis* [8] and *T. politus*; however, the implications for mating behavior continue to be a mystery.

Changes in mating position have been suggested to be associated with many cases of DA in insect genitalia [1,4,11]. Unfortunately we were not able to test this in the case of *T. politus* using live specimens; nevertheless, observations in *Agroeca* Bristowe (1958) and other RTA spiders [12,85] suggest that copulation is achieved by the male climbing over the female and stretching over a side while the female slightly turns her abdomen; this process is alternated between right and left side. In *T. politus*, female genital opening location makes it virtually impossible to have successful mating attempt from a right-side position. Instead, a male must insert both pedipalps always from the left side in relation to the female body. Morphological modifications like: left bulb being slightly bigger (Fig. 5c, f), having a ventrally flattened tegulum (Fig. 5f), and a straight conical conductor (Fig. 5f) instead of the flattened, hook-shaped conductor of the right side (Fig. 5c) are consistent with this hypothesis. In addition, this evidence seems to back the hypothesis discussed by Schilthuizen (2013) and Huber, Sinclair, and Schmitt (2007) stating that in spiders asymmetry is most likely female-initiated and male changes appear as an evolutionary response.

## Conclusions

Genital evolution is a complex and interesting topic. The appearance of asymmetric morphologies is a puzzling phenomenon that has often been overlooked. Here we reported *T. politus* as the first case of directional asymmetry, and the first developmental asymmetry in male genitals in Entelegynae. We also searched for as many cases as possible in taxonomic literature; however, many more might be waiting to be (re)discovered. Our review showed that there have been multiple origins of genital asymmetry in at least thirteen families, and in some cases (e.g. Oonopidae, Pholcidae, Theridiidae, Liocranidae) two or more within the same family. A correct assessment of genital asymmetry based on taxonomic legacy literature is difficult mainly due to the lack of data, description and illustration biases, and number of specimens and variation descriptions.

As has been shown by previous works on genital asymmetry in insects and spiders, there is no single explanation for the evolution of this trait, but some generalizations can be made. In contrast to insects and other arthropod groups, the low number of genital asymmetric species in spiders might indicate that the appearance of these morphological modifications might reduce subsequent speciation rates or even increase extinction rates; specialized lineages tend to have a reduced capacity to diversify and therefore might be considered evolutionary dead ends [86]. However, our observations indicate that cases of sexual asymmetry in spiders, although rare, are more common than was previously thought. Furthermore, they have evolved independently several times but rarely appear isolated and most of the times seem to be clustered within a genus or closely related genera, as in the cases of Oonopidae, Pholcidae, Theridiidae, and probably Liocranidae. The evolution of genital asymmetries in spiders might be a good candidate to be tested as a potential evolutionary dead end.

Several hypotheses for the appearance of asymmetry in spiders have been proposed and include natural selection, sexual selection by female choice and antagonistic co-evolution (not mutually exclusive).We considered *Echinotheridion* and *Tidarren* to be examples of antagonistic co-evolution where the male has evolved self-emasculation in response to the extreme sexual dimorphism in size and aggressive behavior in the female. No selection between left and right is apparent in these genera, thus no directionality is observed. DA cases like *T. politus* seem to support the hypothesis that correlates changes in mating position to genital asymmetry; however, other examples still need to be studied. DA in *T. politus* and some pholcid examples, AS in *Scotinella* and *Asygyna*, and CA cases in *Jacaena*, *Cithaeron* and *Trachelas* support the hypothesis of female-initiated asymmetry in spiders; however, male DA in Oonopidae and AS in some theridiids conflict with this explanation. Further and more detailed study on internal genitalia and comparative study of male right and left pedipalps may yield new and valuable information to explain the evolutionary pattern of genital asymmetry. We hope that this review will aid in the study, development and testing of hypotheses on sexual evolution. We specifically hope it sparks discussions on the complex interactions between males and females, and appearance of interesting phenomena like genital asymmetry.

## Acknowledgements

Thanks to Joe Dulyapat for his great assistance and participation during our fieldwork in Thailand. Thanks to Gustavo Hormiga, Ivan Magalhaes and Martin Ramirez for their remarks and suggestions. Thanks to Fernando Álvarez-Padilla Lab. and Francisco J. Salgueiro-Sepulveda for providing pictures of *Tidarren sisyphoides* and *Triaeris stenaspis*. All *Teutamus* specimens used in this study were collected under permit 5830802 emitted by the Department of National Parks, Wildlife and Plant Conservation, Thailand.

## References

1. Huber BA, Sinclair BJ, Schmitt M. The evolution of asymmetric genitalia in spiders and insects. Vol. 82, Biological Reviews. 2007. p. 647–98.

2. Schilthuizen M. The evolution of chirally dimorphic insect genitalia. Tijdschr voor Entomol [Internet]. 2007;150(December):347–54. Available from: http://www.nev.nl/tve/pdf/te0150347.pdf

3. Schilthuizen M. Something gone awry: unsolved mysteries in the evolution of asymmetric animal genitalia. Anim Biol [Internet]. 2013;63(1):1–20. Available from: http://booksandjournals.brillonline.com/content/journals/10.1163/15707563-00002398

4. Schilthuizen M, de Jong P, van Beek R, Hoogenboom T, zu Schloctern MM. The evolution of asymmetric genitalia in Coleoptera. Philos Trans R Soc B Biol Sci. 2016;371(20150400).

5. van Valen L. A Study of Fluctuating Asymmetry. Evolution (N Y) [Internet]. 1962;16(2):125. Available from: http://www.jstor.org/stable/2406192?origin=crossref

6. Palmer AR. From symmetry to asymmetry: phylogenetic patterns of asymmetry variation in animals and their evolutionary significance. Proc Natl Acad Sci U S A. 1996;93(25):14279–86.

7. van Dongen S. Fluctuating asymmetry and developmental instability in evolutionary biology: past, present and future. J Evol Biol [Internet]. 2006;19(6):1727–43. Available from: http://www.ncbi.nlm.nih.gov/pubmed/17040371

8. Huber BA. Evidence for functional segregation in the directionally asymmetric male genitalia of the spider Metagonia mariguitarensis (Gonzlez-Sponga) (Pholcidae: Araneae). J Zool [Internet]. 2004;262(3):317–26. Available from: http://doi.wiley.com/10.1017/S0952836903004709%5CnCourtMatSP

9. Eberhard WG, Huber BA. Spider Genitalia. Evol Prim Sex Characters Anim. 2010;249–84.

10. Huber BA, Nuñeza OM. Evolution of genital asymmetry, exaggerated eye stalks, and extreme palpal elongation in Panjange spiders (Araneae: Pholcidae). Eur J Taxon [Internet]. 2015;0(169). Available from: http://www.europeanjournaloftaxonomy.eu/index.php/ejt/article/view/291

11. Huber BA. Mating positions and the evolution of asymmetric insect genitalia. Genetica. 2010;138(1):19–25.

12. Foelix RF. Biology of spiders. 3rd ed. Vol. 14, Insect Systematics and Evolution. New York: Oxford University Press; 2011. 419 p.

13. Wheeler WC, Coddington JA, Crowley LM, Dimitrov D, Goloboff PA, Griswold CE, et al. The spider tree of life: phylogeny of Araneae based on target-gene analyses from an extensive taxon sampling. Cladistics. 2016 Dec;

14. Knoflach B, van Harten A. Palpal loss, single palp copulation and obligatory mate consumption in Tidarren cuneolatum (Tullgren, 1910) (Araneae, Theridiidae). J Nat Hist. 2000;34(8):1639–59.

15. Huber BA. Cryptic female exaggeration: The asymmetric female internal genitalia of Kaliana yuruani (Araneae: Pholcidae). J Morphol. 2006;267(6):705–12.

16. Li F, Liu C, Wongprom P, Li S. Sixteen new species of the spider genus Althepus Thorell, 1898 (Araneae: Ochyroceratidae) from Southeast Asia. Zootaxa. 2018;

17. Magalhaes ILF, Brescovit AD, Santos AJ. Phylogeny of Sicariidae spiders (Araneae: Haplogynae), with a monograph on Neotropical Sicarius. Zool J Linn Soc. 2017;

18. Tong Y, Li F, Song Y, Chen H, Li S. Thirty-two new species of the genus Speocera Berland, 1914 (Araneae: Ochyroceratidae) from China, Madagascar and Southeast Asia. Zool Syst. 2019;44(1):1–75.

19. Platnick NI, Dupérré N. The Andean Goblin Spiders of the New Genera Paradysderina and Semidysderina (Araneae, Oonopidae). Bull Am Museum Nat Hist. 2011;364(345):1–121.

20. Lin Y, Pham DS, Li S. Six new spiders from caves of Northern Vietnam (Araneae: Tetrablemmidae: Ochyroceratidae: Telemidae: Symphytognathidae). Raffles Bull Zool. 2009;

21. Huber BA. On American ‘Micromerys’ and Metagonia (Araneae, Pholcidae), with notes on natural history and genital mechanics. Zool Scr. 1996;25(4):341–63.

22. Agnarsson I. Asymmetric female genitalia and other remarkable morphology in a new genus of cobweb spiders (Theridiidae, Araneae) from Madagascar. Biol J Linn Soc. 2006;87(2):211–32.

23. Dondale CD, Redner JH. The insects and arachnids of Canada. Part 9. The sac spiders of Canada and Alaska. Araneae: Clubionidae and Anyphaenidae. In: The insects and arachnids of Canada. 1982. p. 194.

24. Knoflach B. Diversity in the copulatory behaviour of comb-footed spiders (Araneae, Theridiidae). In: Thaler K, editor. Diversität und Biologie von Webspinnen, Skorpionen und anderen Spinnentieren. 2004. p. 161–256.

25. Azevedo GHF, Ott R, Griswold CE, Santos AJ. A taxonomic revision of the ground spiders of the genus Apopyllus (Araneae: Gnaphosidae). Zootaxa. 2016;

26. Opell BD, Beatty JA. The Nearctic Hahniidae (Arachnida: Araneae). Bull Museum Comp Zool. 1976;147(9):393–433.

27. Rivera-Quiroz FA, Alvarez-Padilla F. Three new species of the genus Trachelas (Araneae: Trachelidae) from an oak forest inside the Mesoamerican biodiversity hotspot in Mexico. Zootaxa. 2015;3999(1):95–110.

28. Ruiz GRS, Bonaldo AB. Vagabond but elusive: Two newcomers to the Eastern Amazon (Araneae: Cithaeronidae; Prodidomidae). Zootaxa. 2013.

29. Ahtiainen JJ, Alatalo R V., Mappes J, Vertainen L. Fluctuating asymmetry and sexual performance in the drumming wolf spider Hygrolycosa rubrofasciata. Ann Zool Fennici. 2003;40(40):281–92.

30. Hendrickx F, Maelfait JP, Lens L. Relationship between fluctuating asymmetry and fitness within and between stressed and unstressed populations of the wolf spider Pirata piraticus. J Evol Biol. 2003;16(6):1270–9.

31. Uetz GW, Roberts JA, Wrinn KM, Polak M, Cameron GN. Impact of a catastrophic natural disturbance on fluctuating asymmetry (FA) in a wolf spider. Ecoscience [Internet]. 2009;16(3):379–86. Available from: http://www.bioone.org/doi/abs/10.2980/16-3-3261

32. Uetz GW, McClintock WJ, Miller D, Smith EI, Cook KK. Limb regeneration and subsequent asymmetry in a male secondary sexual character influences sexual selection in wolf spiders. Behav Ecol Sociobiol. 1996;38(4):253–7.

33. Huber BA. Genitalia, fluctuating asymmetry, and patterns of sexual selection in Physocyclus globosus (Araneae: Pholcidae). Rev Suisse Zool. 1996;289–94.

34. Brady AR. The Lynx Spider Genus Oxyopes in Mexico and Central America (Araneae: Oxyopidae). Psyche (New York). 1975;82(2):189–243.

35. Hormiga G. A revision and cladistic analysis of the spider family Pimoidae (Araneoidea: Araneae). Smithson Contrib to Zool. 1994;549:1–104.

36. Kaston BJ. Spider Gynandromorphs and Intersexes. J New York Entomol Soc. 1961;69(4):177–90.

37. Kaston BJ. Abnormal Duplication of the Epigynum and Other Structural, Anomalies in Spiders. Trans Am Microsc Soc. 1963;82(2):220–3.

38. Kaston BJ. Deformities of External Genitalia in Spiders. J New York Entomol Soc. 1963;71(1):30–9.

39. Palmgen P. On the frequency of gynandromorphic spiders. Ann Zool Fennici. 1979;16:183–5.

40. Knoflach B, Van Harten A. The one-palped spider genera Tidarren and Echinotheridion in the Old World (Araneae, Theridiidae), with comparative remarks on Tidarren from America. J Nat Hist. 2006;40(25-26):1483–616.

41. Knoflach B. Copulation and emasculation in Echinotheridion gibberosum (Kulczynski, 1899) (Araneae, Theridiidae). Eur Arachnol. 2002;

42. Penniman AJ. Taxonomic and natural history notes on Phrurolithus fratellus Gertsch (Araneae: Clubionidae). J Arachnol [Internet]. 1978;6:125–32. Available from: http://www.americanarachnology.org/JoA_free/JoA_v6_n2/JoA_v6_p125.pdf

43. Platnick NI, Shadab M. A revision of the bispinosus and bicolor groups of the spider genus trachelas (araneae, clubionidae) in noth and central america and the west Indies. Am Museum Novit. 1974;(2560):34.

44. Platnick NI, Shadab M. A revision of the tranquillus and speciosus groups of the spider genus Trachelas (Araneae, Clubionidae) in North and Central America. Am Museum Novit [Internet]. 1974;(2553):1–34. Available from: http://www.ncbi.nlm.nih.gov/pubmed/23286187

45. Platnick NI, Gajbe UA. Supplementary notes on the ground spider family Cithaeronidae (Araneae, Gnaphosoidea). J Arachnol. 1994;22:32–83.

46. Platnick NI, Dupérré N. The American Goblin Spiders of the New Genus Escaphiella (Araneae, Oonopidae). Bull Am Museum Nat Hist. 2009;328(328):1–151.

47. Deeleman-Reinhold C. Forest spiders of South East Asia: with a revision of the sac and ground spiders (Araneae: Clubionidae, Corinnidae, Liocranidae, Gnaphosidae, Prodidomidae and Trochanterriidae). Leiden: Brill; 2001. 591 p.

48. Dankittipakul P, Tavano M, Singtripop T. Seventeen new species of the spider genus Teutamus Thorell, 1890 from Southeast Asia (Araneae: Liocranidae). J Nat Hist. 2012;46(27-28):1689–730.

49. Dankittipakul P, Tavano M, Singtripop T. Revision of the spider genus Jacaena Thorell, 1897, with descriptions of four new species from Thailand (Araneae: Corinnidae). J Nat Hist. 2013;47(23-24):1539–67.

50. Knoflach B, Benjamin SP. Mating Without Sexual Cannibalism in Tidarren Sisyphoides (Araneae, Theridiidae). J Arachnol. 2003;31(3):445–8.

51. Knoflach B, van Harten A. Tidarren argo sp. nov. (Araneae: Theridiidae) and its exceptional copulatory behaviour: emasculation, male palpal organ as a mating plug and sexual cannibalism. J Zool [Internet]. 2001;254(4):449–59. Available from: http://journals.cambridge.org/abstract_S0952836901000954

52. Agnarsson I. Phylogenetic placement of Echinotheridion (Araneae: Theridiidae) - Do male sexual organ removal, emasculation, and sexual cannibalism in Echinotheridion and Tidarren represent evolutionary replicas? Invertebr Syst. 2006;

53. Alvarez-Padilla F, Hormiga G. A Protocol For Digesting Internal Soft Tissues And Mounting Spiders For Scanning Electron Microscopy. J Arachnol [Internet]. 2007;35(3):538–42. Available from: http://www.bioone.org/doi/abs/10.1636/Sh06-55.1

54. Coddington JA. A Temporary Slide-Mount Allowing Precise Manipulation of Small Structures. Verhandlungen des Naturwissenschaftlichen Vereins Hambg. 1983;26:291–2.

55. WSC. World Spider Catalog Version 20 [Internet]. Natural History Museum Bern. online at http://www.wsc.nmbe.ch [4th april 2019]. 2019. Available from: http://www.wsc.nmbe.ch/

56. Tong Y, Li S. First records of the family Ochyroceratidae (Arachnida: Araneae) from China, with descriptions of a new genus and eight new species. Raffles Bull Zool. 2007;

57. Platnick NI, Dupérré N, Berniker L, Bonaldo AB. The Goblin Spider Genera Prodysderina, Aschnaoonops, and Bidysderina (Araneae, Oonopidae). Bull Am Museum Nat Hist. 2013;(373):1–102.

58. Saaristo MI. Araneae. In: Gerlach J, Marusik Y, editors. Arachnida and Myriapoda of the Seychelles islands. Press Manchester, UK; 2010. p. 306.

59. Platnick NI, Berniker L. The Neotropical Goblin Spiders of the New Genus *Reductoonops* (Araneae, Oonopidae). Am Museum Novit [Internet]. 2014;3811(3811):1–75. Available from: http://www.bioone.org/doi/abs/10.1206/3811.1

60. Platnick NI, Dupérré N, Ubick D, Fannes W. Got Males?: The Enigmatic Goblin Spider Genus Triaeris (Araneae, Oonopidae). Am Museum Novit. 2012;(3756):1–36.

61. Deeleman-Reinhold CL. The Ochyroceratidae of the Indo-Pacific region (Araneae). Raffles bulletin of zoology. 1995.

62. Ferreira RL, Souza MFVR, Machado EO, Brescovit AD. Description of a new Eukoenenia (Palpigradi: Eukoeneniidae) and Metagonia (Araneae: Pholcidae) from Brazilian caves, with notes on their ecological interactions. J Arachnol. 2011;

63. Machado EO, Ferreira RL, Brescovit AD. A new troglomorphic Metagonia Simon 1893 (Araneae, Pholcidae) from Brazil. Zootaxa. 2011;

64. Huber BA, Rheims CA, Brescovit AD. Two new species of litter-dwelling Metagonia spiders (Araneae, Pholcidae) document both rapid and slow genital evolution. Acta Zool. 2005;

65. González AP, Huber BA. Metagonia debrasi n. sp. the first species of the genus Metagonia Simon in Cuba (Pholcidae, Araneae). Rev Arachnol. 1999;

66. Gertsch WJ, Ennik F. The spider genus Loxosceles in North America, Central America, and the West Indies (Araneae, Loxoscelidae). Bull Am Museum Nat Hist. 1983;175:264–360.

67. Lotz LN. An update on the spider genus Loxosceles (Araneae: Sicariidae) in the Afrotropical region, with description of seven new species. Zootaxa. 2017;4341(4):475–94.

68. Wang C, Li S. Three new species of Telemidae (Araneae) from Western Africa. Zootaxa. 2011;

69. Dupérré N, Tapia E. Discovery of the first telemid spider (Araneae, Telemidae) from South America, and the first member of the family bearing a stridulatory organ. Zootaxa. 2015;

70. Wang C, Li S. Four new species of the spider genus Telema (Araneae, Telemidae) from Southeast Asia. Zootaxa. 2010;

71. Wang C, Li S. New species of the spider genus Telema (Araneae, Telemidae) from caves in Guangxi, China. Zootaxa. 2010;

72. Lin Y, Li S. Long-legged cave spiders (Araneae, Telemidae) from Yunnan-Guizhou Plateau, southwestern China. Zootaxa. 2010;

73. Platnick NI. A revision of the ground spider family Cithaeronidae (Araneae, Gnaphosoidea). Am Museum Novit. 1991;3018:1–13.

74. Platnick NI, Shadab M, Sorkin L. On the Chilean Spiders of the Family Prodidomidae (Araneae, Gnaphosoidea), with a Revision of the Genus Moreno Mello-Leitão. Am Museum Novit. 2005;1–31.

75. Gajbe UA. Araneae: Arachnida. In: Fauna of Madhya Pradesh (including Chhattisgarh), State Fauna Series. Zool Surv India, Kolkata. 2007;15(1):419–540.

76. Holwell GI, Herberstein ME. Chirally dimorphic male genitalia in praying mantids (Ciulfina: Liturgusidae). J Morphol. 2010;

77. Gertsch WJ. The spider genus Loxosceles in South America (Araneae, Scytodidae). Bull Am Museum Nat Hist. 1967;136:117–74.

78. Wang C, Li S. Four new species of the subfamily Psilodercinae (Araneae: Ochyroceratidae) from Southwest China. Zootaxa. 2013;

79. Swaddle JP, Witter MS, Cuthill IC. The analysis of fluctuating asymmetry. Animal Behaviour. 1994. p. 986–9.

80. Palmer AR, Strobeck C. Fluctuating asymmetry and developmental stability: heritability of observable variation vs. heritability of inferred cause. J Evol Biol. 1997;

81. Schilthuizen M, Davison A. The convoluted evolution of snail chirality. Naturwissenschaften. 2005.

82. Holwell GI, Kazakova O, Evans F, O’Hanlon JC, Barry KL. The functional significance of chiral genitalia: Patterns of asymmetry, functional morphology and mating success in the praying mantis Ciulfina baldersoni. PLoS One. 2015;10(6).

83. Palmer AR. Symmetry breaking and the evolution of development. Science. 2004.

84. Bristowe WS. The world of spiders. The world of spiders. London: Collins; 1958. 304 p.

85. Brady AR. The lynx spiders of North America, north of Mexico (Araneae: Oxyopidae). Bull Museum Comp Zool. 1964;131:429–518.

86. Agnarsson I, Avilés L, Coddington JA, Maddison WP. Sociality in theridiid spiders: repeated origins of an evolutionary dead end. Evolution (N Y). 2006;

